# Quantifying Interpretation Reproducibility in Vision Transformer Models with TAVAC

**DOI:** 10.1101/2024.01.18.576252

**Authors:** Yue Zhao, Dylan Agyemang, Yang Liu, Matt Mahoney, Sheng Li

## Abstract

The use of deep learning algorithms to extract meaningful diagnostic features from biomedical images holds the promise to improve patient care given the expansion of digital pathology. Among these deep learning models, Vision Transformer (ViT) models have been demonstrated to capture long-range spatial relationships with more robust prediction power for image classification tasks than regular convolutional neural network (CNN) models, and also better model interpretability. Model interpretation is important for understanding and elucidating how a deep learning model makes predictions, especially for developing transparent models for digital pathology. However, like other deep learning algorithms, with limited annotated biomedical imaging datasets, ViT models are prone to poor performance due to overfitting, which can lead to false predictions due to random noise. Overfitting affects model interpretation when predictions are made out of random noise. To address this issue, we introduce a novel metric – Training Attention and Validation Attention Consistency (TAVAC) – for evaluating ViT model degree of overfitting on imaging datasets and quantifying the reproducibility of interpretation. Specifically, the model interpretation is performed by comparing the high-attention regions in the image between training and testing. We test the method on four publicly available image classification datasets and two independent breast cancer histological image datasets. All overfitted models exhibited significantly lower TAVAC scores than the good-fit models. The TAVAC score quantitatively measures the level of generalization of model interpretation on a fine-grained level for small groups of cells in each H&E image, which cannot be provided by traditional performance evaluation metrics like prediction accuracy. Furthermore, the application of TAVAC extends beyond medical diagnostic AI models; it enhances the monitoring of model interpretative reproducibility at pixel-resolution in basic research, to reveal critical spatial patterns and cellular structures essential to understanding biological processes and disease mechanisms. TAVAC sets a new standard for evaluating the performance of deep learning model interpretation and provides a method for determining the significance of high-attention regions detected from the attention map of the biomedical images.

## Main

Application of deep learning in the analysis of biomedical histological images is rapidly advancing the field of digital pathology by providing nuanced pattern recognition that surpasses traditional methods (Lo & Hung, 2022; Hu, et al., 2022; Hu, et al., 2021; Yang & Stamp, 2021). Deep learning models, e.g., convolutional neural networks (CNN), have been successfully applied to histopathology images for predictive purposes such as classifying tumor (Litjens, et al., 2018), predicting mutation (Coudray, et al., 2018), and identifying cancer subtypes (Khosravi, et al., 2018). The clear advantage of deep learning algorithms—particularly when trained on vast and diverse datasets—is their potential to offer unprecedented accuracy in identifying and classifying cellular structures and abnormalities, which is critical for early and precise disease diagnosis. However, key challenges include the scarcity of high-quality, labeled histological datasets for training, high computational demands, and the necessity for transparent models that digital pathologists can trust. Nevertheless, the potential benefits of deep learning, such as automating time-consuming diagnostic tasks and unveiling new histological features correlated with patient outcomes, are driving research and innovation forward.

Through training on large histological datasets, deep learning models acquire the ability to extract relevant features that give insight into tumor regions and biomarkers for cancer classification. Among deep learning algorithms, the Vision Transformer (ViT) model has exhibited strong prediction power in image classification tasks (Raghu, et al., 2021). Unlike conventional neural networks (CNN), ViT models utilize self-attention techniques to robustly capture long-range relationships that CNNs cannot (Mao, et al., 2022). ViT models process images by splitting them into patches, embedding them into a lower-dimensional space, and adding positional encodings and a class token. These sequences pass through a transformer encoder and the class token’s values are used in a multilayer perceptron (MLP) to classify the image (Islam, 2022). Importantly, in the ViT architecture, the attention mechanism dynamically computes weights for input tokens based on their contextual relevance to a designated query. These computed weights serve to determine the influence of each individual token on the resultant representation, facilitating the model’s ability to discern intricate relationships and contextual nuances among tokens.

Transparent, explainable, and interpretable are pivotal characteristics of trustworthy AI systems (Zhang, et al., 2019). Like other deep learning models, a major limitation of ViT models is their tendency to overfit when data size is limited, which is a common problem for biomedical imaging datasets with annotations, e.g., datasets generated using histological methods, computed tomography, X-ray, ultrasound, and optical coherence tomography. Overfitting in machine learning models manifests as an excessive adaptation to the training data, incorporating noise and specific data idiosyncrasies, which impairs the model’s ability to generalize to new data (Dietterich, 1995; Lever, et al., 2016; Ying, 2019). This overfitting is often attributable to the noise inherent in a non-representative training dataset and an overly complex hypothesis that causes the model to prioritize variance over bias, leading to inconsistent predictive performance. A model that is optimized for generalization effectively discerns relevant patterns from noise in the training data and maintains a balance between hypothesis complexity and predictive consistency (Ying, 2019). The majority of prior studies on overfitting focused on the predictive performance of deep learning models. At the same time, the generalization of deep learning model interpretation through attention visualization has been understudied. Ramampiandra & Chollet previously commented that overfitted models impede the interpretation of machine learning models (Ramampiandra & Chollet, 2023), yet no solution is proposed to solve the problem.

Model interpretation for ViT models is carried out through analysis of attention layers, which are the weight on the input representing the focus of the model (Dosovitskiy, et al., 2020; Abnar & Zuidema, 2020). Given the increasing interest in leveraging attention maps to explain predictive models, it is critical to assess how overfitting impacts the model interpretability. Ideally, the attention maps should highlight semantically meaningful features in the image, so that similar input images will have attention highlighting similar features. Generalization of the model interpretation, when the model encounters various datasets, needs to be controlled.

Unlike common images, biomedical images (e.g., histological staining) are highly vulnerable to overfitting due to its obscure characteristics, as one cannot explicitly decipher if the prediction is based on a true biologically meaningful pattern or random noise. Overfitting has particularly serious implications in the analysis of histological image data. Incorrect or misleading predictions can have detrimental effects on clinical diagnoses in the healthcare sector (Lo & Hung, 2022; Hu, et al., 2022; Hu, et al., 2021; Yang & Stamp, 2021). Because of this, assessing the degree of self-consistency in ViT model interpretations is essential to ensure transparent models, accurate predictions, and robust conclusions. Outside of deep learning, this issue has been confronted before in the literature on functional brain imaging, where high-dimensional statistical models are fit to brain scans and scientific inferences are made based on the model parameters (Strother, et al., 2002). In that setting, it was found that model self-consistency and predictive performance do not always align exactly; the highest-performing models imply biased scientific inferences. The solution is to explicitly test for self-consistency by comparing fit model parameters on different splits of the data. Building on this idea and applying it to attention maps, we have developed Training Attention and Validation Attention Consistency (TAVAC) that evaluates the performance of model interpretation by quantifying the stability of the attention maps between when an image is used for training or validation during model fitting. While the other evaluation metrics (accuracy, F-1 score, mean squared error, etc.) address whether accurate predictions are made, our novel algorithm of Training Attention and Validation Attention Consistency (TAVAC) evaluates the generalization of model interpretation.

## Results

### Overview of TAVAC algorithm to delineate the generalization of model interpretation

To address the impact of overfitting on ViT model interpretation of biomedical imaging data, we sought to develop a metric to evaluate model robustness in the context of attention-driven model behaviors, as depicted in Fig 1. We took inspiration from the established K-fold cross-validation technique, with a particular focus on a two-fold implementation (Strother, et al., 2002; Yadav & Shukla, 2016). The dataset D is bifurcated into two distinct folds, employing one for training and the other for validation purposes in the initial stage (stage 1). Specifically, subsequent to the training of a ViT model (Dosovitskiy, et al., 2020) utilizing the first training subset, its performance is evaluated against the validation subset. In the ensuing stage (stage 2), the roles of the subsets reverse, wherein the initial training dataset is deployed for validation and the initial validation dataset is deployed for training, followed by a reiteration of the training and evaluation process. In stage 2, the ViT has to be reinitialized. TAVAC algorithm examines the model’s internal consistency by juxtaposing the *pixel-level* attention maps for images from training in stage 1 against validation in stage 2. Within each stage, attention maps for individual images are formulated by employing attention rollout methodology (Abnar & Zuidema, 2020). Specifically, when an image is designated as ‘training data,’ the resultant attention map is categorically defined as Training Attention; and conversely, when designated as ‘validation data,’ it is defined as Validation Attention. The TAVAC score is subsequently articulated as the consistency between Training Attention and Validation Attention, measured using Pearson correlation.

**Figure 1:**
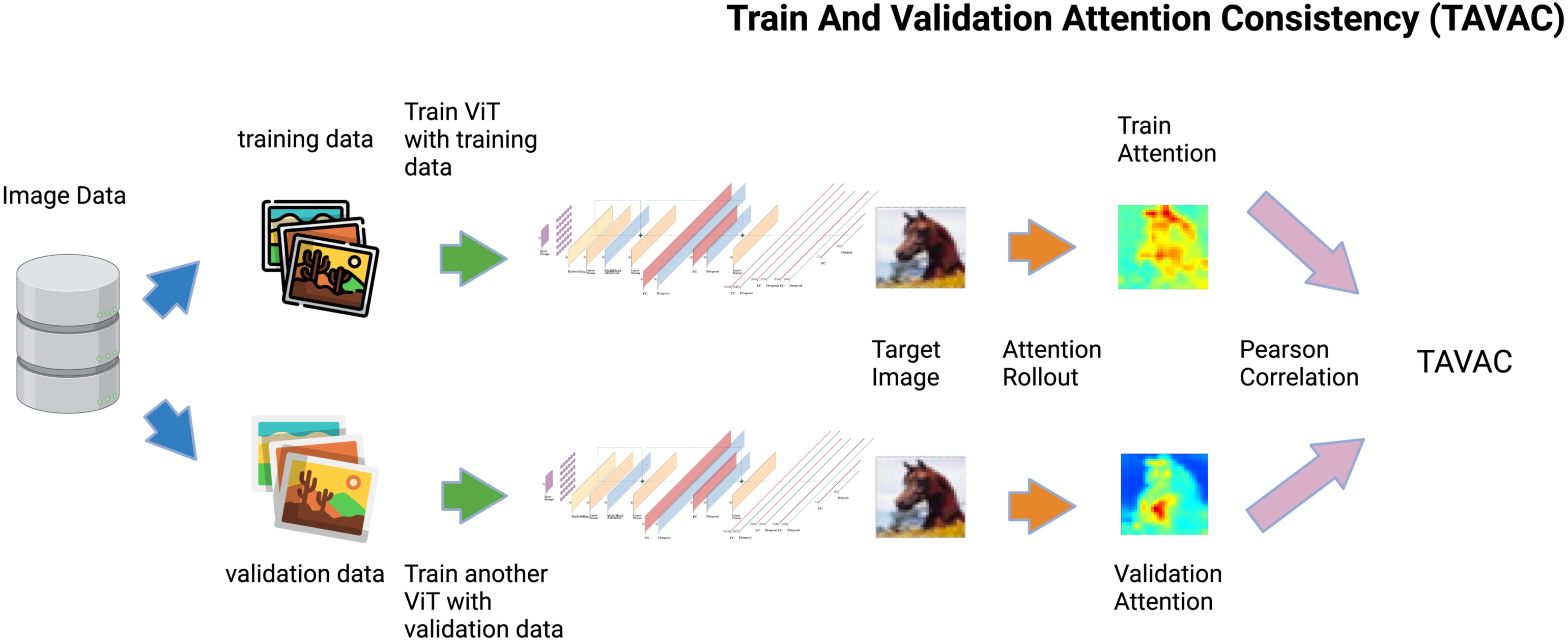
General workflow of the Train and Validation Attention Consistency (TAVAC) algorithm. The TAVAC Algorithm quantifies the generality of ViT Model Interpretation. The schematic outlines the two-stage process where training and validation subsets are used alternately to develop a ViT model and assess the internal consistency of model interpretation. Attention maps are generated via attention rollout for the same image (horse image from the training subset in this example) from ViT models trained with either training or validation subsets, with TAVAC scores computed based on the Pearson correlation between these pixel-level attention map distributions, reflecting model attention consistency across different ViT models over the same horse image.

### Evaluate TAVAC using four image benchmark datasets as proof of concept

To evaluate the TAVAC algorithm, we trained two types of pretrained ViT models by varying epochs settings using transfer learning – a good-fit model and an overfitted model – and examined their attention maps and TAVAC score distribution (Table 1). We evaluated TAVAC algorithm using four well-known image classification benchmarking datasets: CIFAR-10, MNIST, and Food-101, and Cats vs. Dogs (Krizhevsky, 2009; Lecun, et al., 2010; Bossard, et al., 2014; Golle, 2008). The various image types and groups of classes among these datasets allow us to prove TAVAC effectiveness and generalization: (1) The CIFAR-10 dataset consists of 60,000 of 32 × 32 color images in 10 classes, with 6000 images attributed to each class. (2) The MNIST dataset consists of 70,000 28 × 28 grayscale images of ten classes of handwritten digits, i.e., numerals. (3) The Food-101 dataset consists of 101,000 images containing various foods of 101 classes with varying dimensions. (4) Lastly, the Cats vs. Dogs dataset, a subset of the Animals Species Image Recognition for Restricting Access (Asirra) dataset, contains 23,442 cat and dog images of varying dimensions. As expected, excessively high epoch settings led to an overfitted model while comparatively lower epoch settings generated good-fit models with comparable training and validation accuracies (Table 2), consistent with general machine learning knowledge (Lecun, et al., 2012). The settings corresponding to good-fit and overfitted models varied for each dataset, with overfitting on MNIST and CATS VS. DOGS requiring substantial epoch numbers, while overfitting on CIFAR-10 and Food-101 required much lower epoch numbers (Table 1). We assessed TAVAC’s ability to distinguish the impact of overfitting on model interpretation using each of the datasets as described below to show that the overfitted and good-fit ViT models have different TAVAC score distributions.

**Table 1.** Hyperparameters for training good-fit and overfitted ViT models.

| Dataset | Types | Number of Epochs | Learning Rate | Weight Decay | Batch Size |
| --- | --- | --- | --- | --- | --- |
| CIFAR10 | Good-fit | 10 | 2e-5 | 0.01 | 10 |
|  | Overfitted | 100 |  |  |  |
| MNIST | Good-fit | 1500 | 1e-4 | 0 | 4 |
|  | Overfitted | 2000 |  |  |  |
| FOOD101 | Good-fit | 10 | 1e-4 | 0 | 64 |
|  | Overfitted | 100 |  |  |  |
| CATS VS. DOGS | Good-fit | 1000 | 1e-4 | 0 | 32 |
|  | Overfitted | 5000 |  |  |  |

**Table 2.** Training and validation accuracies of ViT models.

| Dataset | Model Type | Stage 1 Training Accuracy | Stage 1 Validation Accuracy | Stage 2 Training Accuracy | Stage 2 Validation Accuracy |
| --- | --- | --- | --- | --- | --- |
| CIFAR10 | Good-fit | 0.90 | 0.93 | 0.968 | 0.942 |
|  | Overfitted | 0.944 | 0.74 | 0.982 | 0.752 |
| MNIST | Good-fit | 0.958 | 0.958 | 0.96 | 0.966 |
|  | Overfitted | 0.962 | 0.942 | 0.964 | 0.946 |
| FOOD101 | Good-fit | 0.982 | 0.946 | 0.952 | 0.939 |
|  | Overfitted | 0.987 | 0.915 | 0.988 | 0.914 |
| CATS VS. DOGS | Good-fit | 1.0 | 0.982 | 0.972 | 1.0 |
|  | Overfitted | 1.0 | 0.962 | 0.998 | 0.958 |

### TAVAC delineates the generalization of model interpretation by image benchmark datasets

We applied TAVAC to good-fit and overfitted ViT models trained using 1000 randomly selected images from the CIFAR-10 dataset (500 for each fold). We generated a good-fit model using 10 epochs based on a pre-trained ViT model (Dosovitskiy, et al., 2020), which has comparable training and validation accuracies, in both stage 1 and stage 2 (Table 2). We then compared each image’s training attention from stage 1 and validation attention from stage 2 (and vice versa, Fig. 1) to determine the TAVAC score distribution for each model. As expected with the good-fit model, both training attention and validation attention have high attention regions (HAR) primarily focused on the horse object (Fig.2a). Next, we examined the TAVAC of an overfitted model of 100 epochs for the same dataset. This variation of the model possessed a lower validation accuracy than training accuracy, in both stage 1 and stage 2 (Table.2). Moreover, the overfitted model’s HAR of training attention focused on the horse object, while the HAR of validation attention fixated on a non-meaningful section of the image, away from the object of interest (Fig.2b).

We then used a scatterplot to quantify the TAVAC at the pixel level. Using the horse image as an example (Fig. 2a), we plotted *pixel-level* training attention against validation attention for both the good-fit and overfitted models. The good-fit model’s scatterplots (Fig.2a third column) exhibited a strong positive correlation between Training Attention and Validation Attention, yielding a Pearson correlation coefficient (r-value) of 0.95, which we denote as the TAVAC score. In contrast, the overfitted model’s scatterplot (Fig.2b) exhibited an extremely poor correlation between training attention and validation attention, and the r-value is 0.37. As a holistic analysis, we employed TAVAC on every image and compare the distribution between good-fit and overfitted models (Fig. 2c). Indeed, the TAVAC score of overfitted model (median: 0.644, inner quartile range IQR: 0.327) is significantly lower and more variable than good-fit model (median: 0.960, IQR: 0.045, P-value < 0.001, Kolmogorov Smirnov KS test), thus, the TAVAC scores declines substantially when the ViT model is overfitted for the CIFAR-10 dataset.

**Figure 2:**
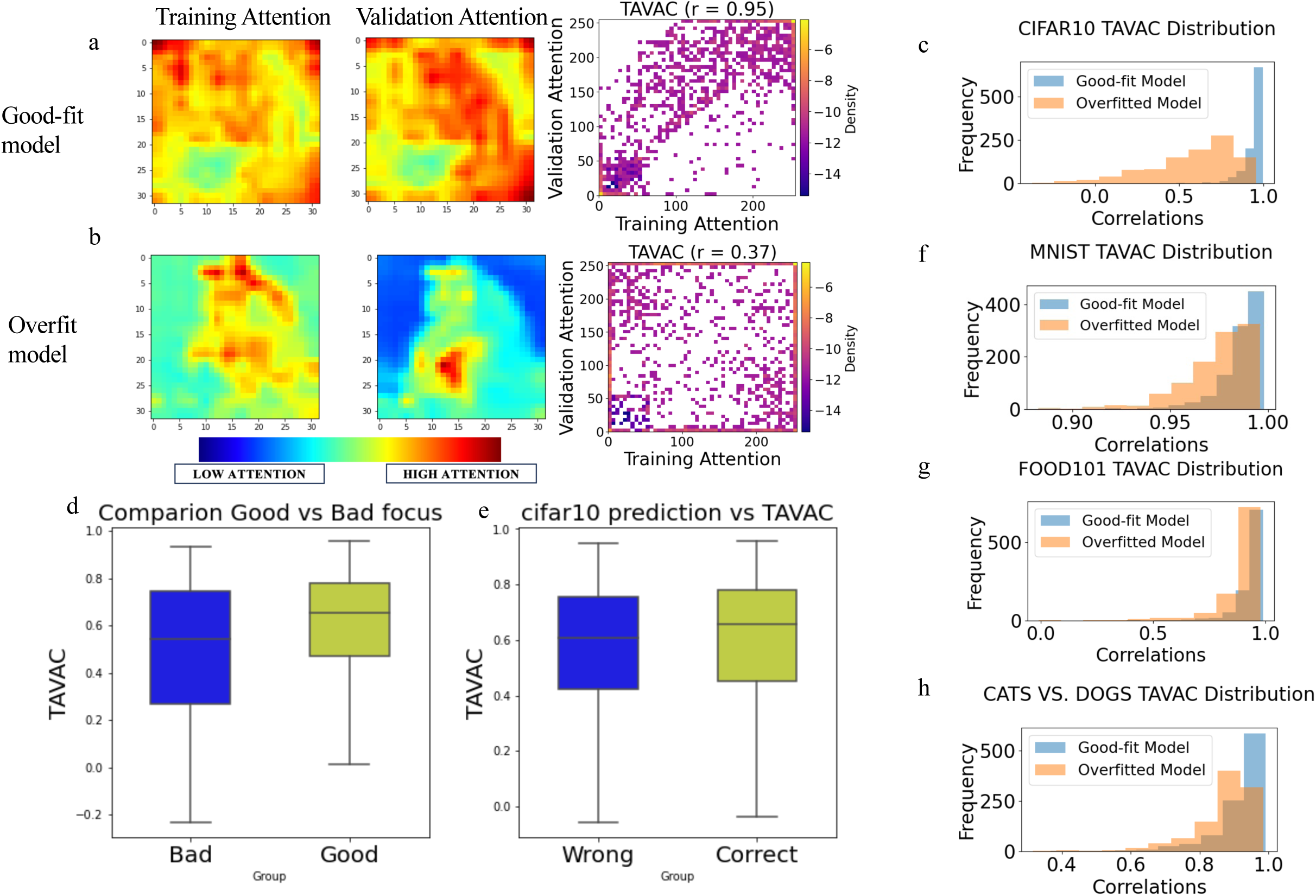
TAVAC Evaluation on Image Datasets Using Good-Fit vs. Overfitted ViT Models. (a-b) Pixel-Level TAVAC Evaluation for CIFAR10 Horse Image Analysis: TAVAC scores are depicted for a horse image, with (a) well-fitted models showing strong attention correlation (r = 0.95) and (b) overfitted models displaying attention divergence (r = 0.37). Attention maps use red for high attention, blue for low, and green for intermediate. (c) TAVAC Score Distributions: Histograms for CIFAR10 illustrate the frequency distribution of TAVAC scores, blue for well-fitted and yellow for overfitted, highlighting diminished attention consistency with increased overfitting. (d-e) Focus and Prediction Accuracy Comparisons: Box plots compare TAVAC scores for images with (d) ‘Good Focus’ versus ‘Bad Focus’ and (e) ‘Correct’ versus ‘Wrong’ predictions. ‘Good Focus’: manually annotated images where high attention is maintained on the object of interest during validation in well-fitted cases; ‘Bad Focus’: manually annotated images with poor attention focus during validation in overfitted cases. ‘Correct’ and ‘Wrong’ depict images that the ViT model predicted accurately or inaccurately, respectively. (f-h) Cross-Dataset Validation: TAVAC distributions for MNIST, Food-101, and Cats vs. Dogs validate the CIFAR10 findings, with consistent attention reduction in overfitted models.

To evaluate the robustness of TAVAC in identifying meaningful high-attention regions, we visually inspected all the 1000 images. We annotated 114 images, in which HAR of the validation attention of the *overfitted* model does not overlap with the object of interest and is defined as ‘bad focus’ group (Fig. S1). The remaining images are annotated as ‘good focus’ group. The bad focus group exhibits significantly lower TAVAC scores than good focus group (Fig. 2d, t-test, *P*-value = 0.00001). Notably, the TAVAC scores of the bad focus group are as low as −0.2, while the scores of good focus group are all positive. This suggests that TAVAC can accurately identify instances where the ViT model doesn’t utilize the meaningful image features for prediction by attributing a low score to them. Moreover, we compared the TAVAC scores of incorrectly predicted images, labeled as wrong, and correctly predicted images, labeled as correct. After our analysis, we conclude that incorrectly predicted images have lower TAVAC scores than correctly predicted images (Fig. 2e, *P*-value = 0.025, t-test). However, the two groups show sizeable overlap, indicating that the prediction performance is not identical to the TAVAC score, and TAVAC can serve as an independent evaluator for ViT model interpretation. We further applied the TAVAC algorithm to three additional datasets: MNIST, FOOD101, and Cats vs. Dogs. A summary of the model’s performance is provided in Table 1. In the MNIST dataset, the overfitted model (median: 0.979) exhibited modestly lower TAVAC scores than the good fit model (median: 0.989) with a statistical significance (Fig. 2f, *P*-value = 4.68 × 10^-65^, KS test). While the overfitted model (median: 0.919) of the FOOD101 dataset exhibited lower TAVAC scores than the good fit model (median: 0.955) with a statistical significance (Fig. 2g, *P*-value = 1.21 × 10^-62^, KS test). Similarly, for the Cats vs. Dogs dataset, the overfitted model (median: 0.895) showed lower TAVAC scores than the good fit model with a statistical significance (median: 0. 942, Fig. 2h, *P*-value = 1.23 × 10^-57^, KS test). Similar to the CIFAR-10 dataset, the manually annotated bad focus group in Cats vs. Dogs dataset exhibits significantly lower than TAVAC scores than good focus group (Fig. S2, t-test, *P*-value = 0.018). Overall, the TAVAC algorithm shows robustness in distinguishing the inaccurate attention maps of overfitted models from accurate attention maps in good-fit models, across various types of images, e.g., hand-writing number “8”, hamburger, and a dog (Fig. 3). Overall, as overfitting increased through varying epoch numbers, TAVAC scores decreases in all four imaging benchmarking datasets. The results suggest that TAVAC robustly quantifies the impact of overfitting on ViT model interpretation and the attention maps, which contribute to the transparency of deep learning models.

**Figure 3:**
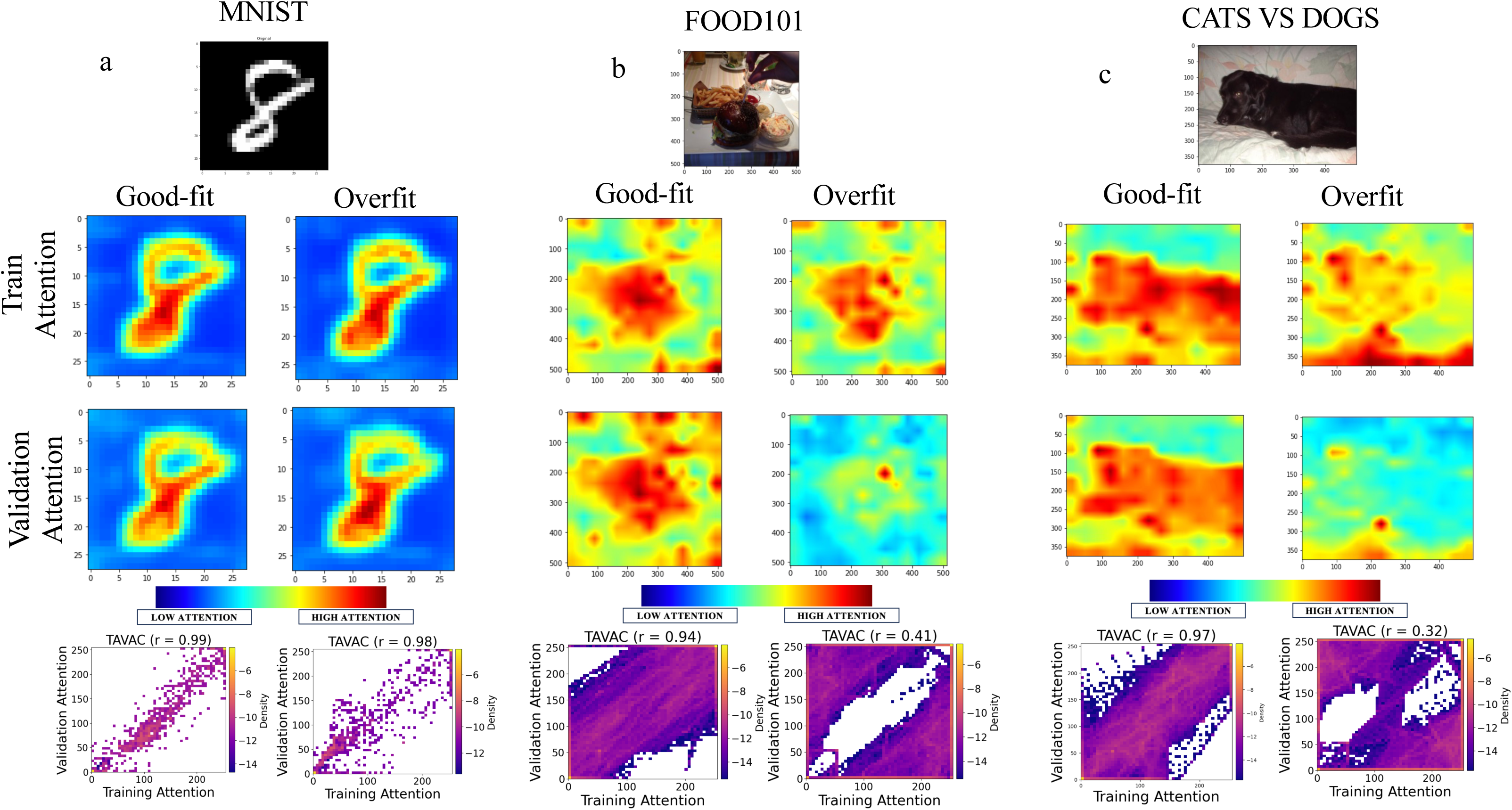
Pixel-Level TAVAC Analysis for MNIST, Food-101, and Cats vs. Dogs. (a) MNIST ‘8’ Image: TAVAC analysis reveals high attention consistency for a well-fitted model (TAVAC score barely changes with slight overfitting). (b) Food-101 Hamburger Image: Well-fitted model attention maps closely align (high TAVAC score of 0.94), whereas overfitted model shows significant attention misalignment (low TAVAC score of 0.41). (c) Cats vs. Dogs Dog Image: Attention is consistent in well-fitted models (TAVAC score of 0.94), with a substantial drop in overfitted models (TAVAC score of 0.32). Density scatterplots accompany each analysis, with purple indicating high density (consistent attention) and yellow indicating low density (inconsistent attention). These comparisons demonstrate that TAVAC scores decrease as model overfitting increases.

### Application to two independent breast cancer pathological image datasets

To examine the overfitting impact on ViT model interpretation on biomedical images, we next applied the TAVAC method to the transfer learning models trained using two independent breast cancer pathological image datasets. The first breast cancer H&E dataset (He, et al., 2020) consists of 23 patients, 68 tissue sections, and 30,612 spots (diameter: 100 µm). For each patient, we used a collection of 3 microscopic slide images of high-resolution H&E-stained tissue, all of which have tumor or non-tumor annotation by pathologists. We subsequently applied TAVAC algorithm by dividing the data set into two random partitions, each comprised of approximately 15,000 spots. We then trained the pretrained ViT model (Dosovitskiy, et al., 2020) to predict tumor and non-tumor classes using H&E image spots (example in Fig. 4a) via transfer learning (Table 3-4). The confusion matrix of the ViT model prediction is shown in Fig. 4b. We derived training and validation attention maps using 1000 randomly selected H&E image spots, similar to the procedure for the image benchmark datasets. The ViT model of the breast cancer H&E image exhibits high TAVAC scores for the majority of tissue spots with minimal spread (Fig. 4c, median: 0.933, IQR: 0.070). We further visualized the consistency of the training and validation attention of H&E images with various TAVAC scores. The high-TAVAC-score images (Fig. 4d and Fig. S3) exhibit high attention regions (HARs) of both training and validation attention enriched for tumor and cellular regions (dark color). While the low-TAVAC-score images have HARs of validation attention that are inconsistent with the training attention, suggesting non-specific image features were used for the prediction of tumors (Fig. 4e and Fig. S4). All predictions for training and validation are correct and all samples in Fig.4d-e are tumor samples. This is consistent with expectation, as the ViT is trained to predict the presence of tumor tissue, thus the HARs are expected to overlap with the tumor regions in the H&E images. This suggests that the high-TAVAC score image HARs are more biologically meaningful than the low-TAVAC score image HARs. Thus, the results showed that the TAVAC score can serve as a quality control metric of ViT model interpretation to rank H&E images for attention maps of biologically meaningful HAR.

**Figure 4.**
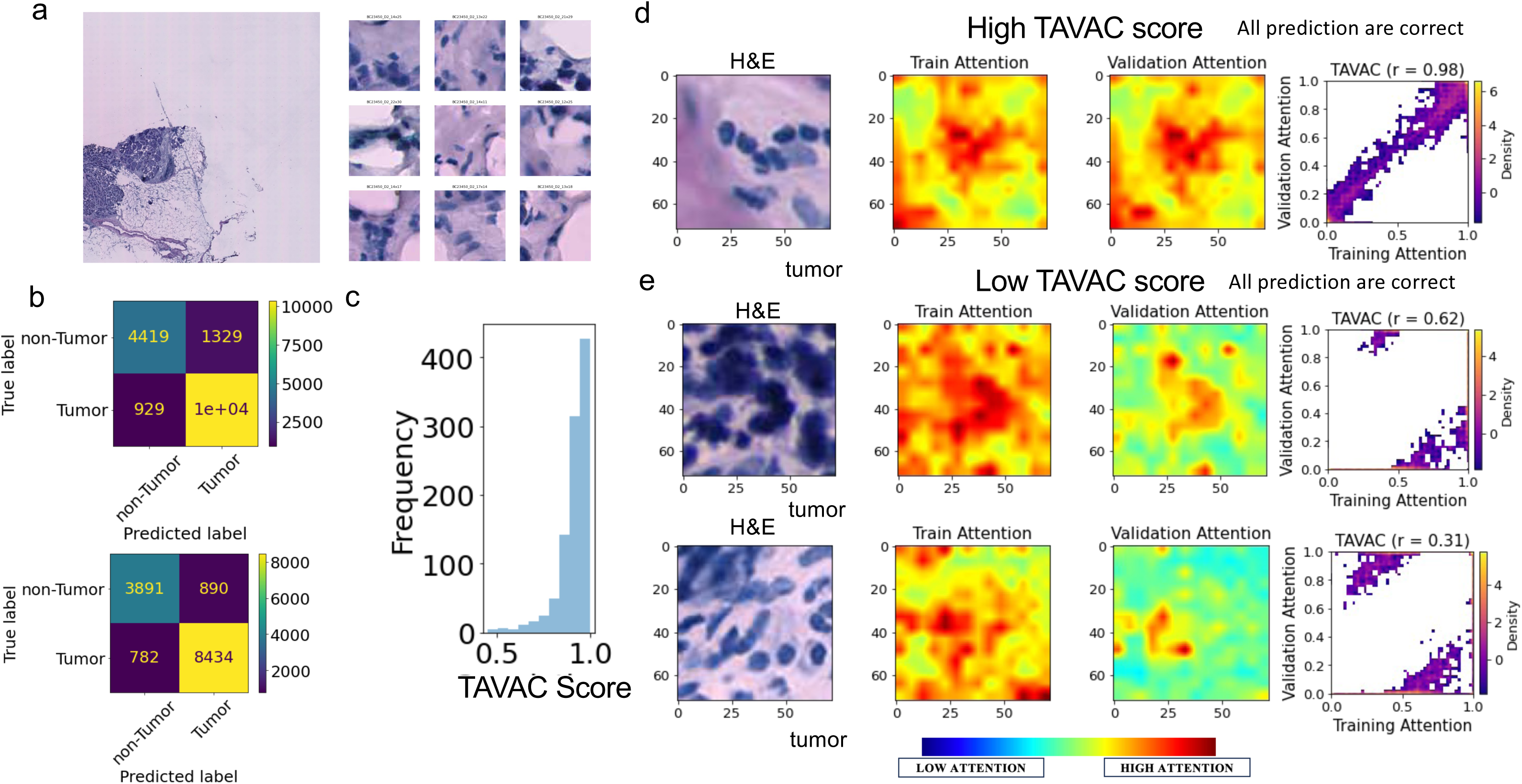
TAVAC Quantifies the Slide-and Pixel-level Generality of ViT model Interpretation using Breast Cancer Pathology Images. (a) Displays paired H&E images of breast tissue, with a full tissue cut on the left and a magnified spot image on the right. (b) Presents confusion matrices for validation data: the TAVAC stage 1 model at the top and the TAVAC stage 2 model below. (c) Histogram of TAVAC scores shows consistent training-validation attention across the majority of spots. (d) A high TAVAC score (0.96) case where training and validation attentions align on tumor spots, as shown in the density scatterplot. (e) A low TAVAC score cases (0.62 and 0.31) displays a marked difference in attention between training (accurately focused on tumor regions) and validation (misaligned). Both (d) and (e) confirm tumor presence with correct model predictions across training and validation.

**Table 3.** Hyperparameters for training breast cancer pathological image ViT models.

| Dataset | Number of Epochs | Learning Rate | Weight Decay | Batch Size |
| --- | --- | --- | --- | --- |
| Breast cancer | 500 | 1e-5 | 0 | 32 |
| HER-2 | 500 | 1e-5 | 0.001 | 10 |

**Table 4.** Training and validation accuracies of breast cancer ViT models.

| Dataset | Stage 1<br>Training<br>Accuracy | Stage 1<br>Validation<br>Accuracy | Stage 2<br>Training<br>Accuracy | Stage 2<br>Validation<br>Accuracy |
| --- | --- | --- | --- | --- |
| Breast<br>cancer | 0.994 | 0.867 | 0.993 | 0.880 |
| HER-2 | 0.995 | 0.847 | 0.991 | 0.846 |

For independent verification, we performed the TAVAC analysis on another H&E image dataset: HER2 breast cancer dataset (Andersson, et al., 2021) from eight HER2-positive patients. From each patient’s respective tumor specimen, three adjacent or six evenly spaced tissue sections were extracted (Andersson, et al., 2021). Additionally, a pathologist annotated a representative tissue section from each distinct tumor according to the morphology exhibited by the H&E image. The annotated data has nine replicates of H&E images totaling around 3800 spots. The categorizations include *in situ* cancer, invasive cancer, adipose tissue, immune infiltrate, or connective tissue (Andersson, et al., 2021). We used invasive cancer or non-invasive cancer as class labels for each spot. We divided the spot H&E image data set into two random partitions to train ViT models (Table 3-4) for invasive cancer prediction (example in Fig. 5a). The confusion of classification is shown in Fig. 5b. After randomly selecting a subset of 500 spot images from each partition, we generated their respective attention maps. The ViT model of the HER-2 breast cancer H&E image exhibits high TAVAC scores for the majority of tissue spots with minimal spread (Fig. 5c, median: 0.971, IQR: 0.031). We then compared the training and validation attention of H&E images with various TAVAC scores. The high-TAVAC-score image (Fig. 5d) exhibits high attention regions (HARs) of both training and validation attention enriched for tumor and cellular regions (dark color). Conversely, the low-TAVAC-score images have HARs of validation attention that are inconsistent with the training attention, where the HARs of validation attention are enriched for non-tumor regions (Fig. 5e). All predictions for training and validation are correct. Figure 5d and Figure 5e bottom are invasive cancer samples while Figure 5e top represents a non-invasive-cancer sample. The results collectively confirmed the TAVAC score as a quality control metric of ViT model interpretation to rank H&E images with attention maps of biologically meaningful HAR. Thus, we visualized the high vs. low TAVAC score for each spot on the H&E image (Fig. 6a and S5). to identify the spots to filter out for interpretation study. The low-TAVAC-score spots that should be filtered out (< 0.9) are shown as black dots (Fig. 6b) given the high discrepancy between training and validation attention maps. Moreover, the low TAVAC spots are spread across the tissue without association with specific pathological categories (Fig. 6c and S5).

**Figure 5.**
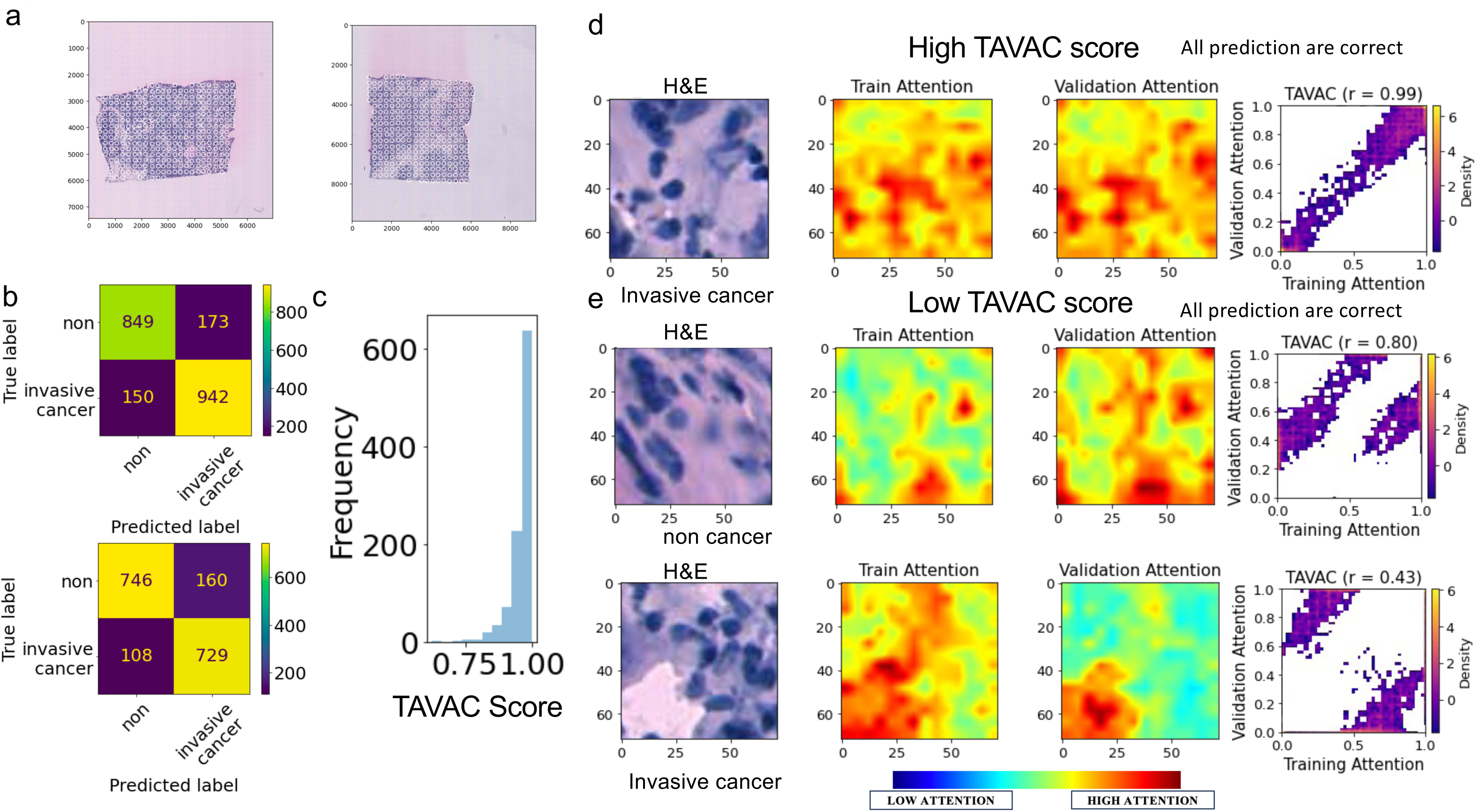
Independent Validation of TAVAC performance using HER2+ Breast Cancer Pathology Image ViT Model Interpretation. (a) Displays a sample H&E stained full tissue cut and a corresponding detailed spot image of breast tissue. (b) Shows confusion matrices for HER2+ breast cancer two-stage models: stage 1 (top) and stage 2 (bottom) model validation. (c) Illustrates a histogram of TAVAC scores, indicating consistent attention between training and validation for the majority of evaluated spots. (d) High TAVAC score (0.96) example in non-cancer H&E images where training and validation attention converge on identical regions, supported by a density scatterplot. (e) Low TAVAC score (0.63) case in invasive cancer H&E images, exhibiting a clear visual discrepancy in attention focus between training and validation, especially on dark tumor spots, as shown in the density scatterplot. For both (d) and (e), the model predictions are accurate in both the training and validation phases.

**Figure 6.**
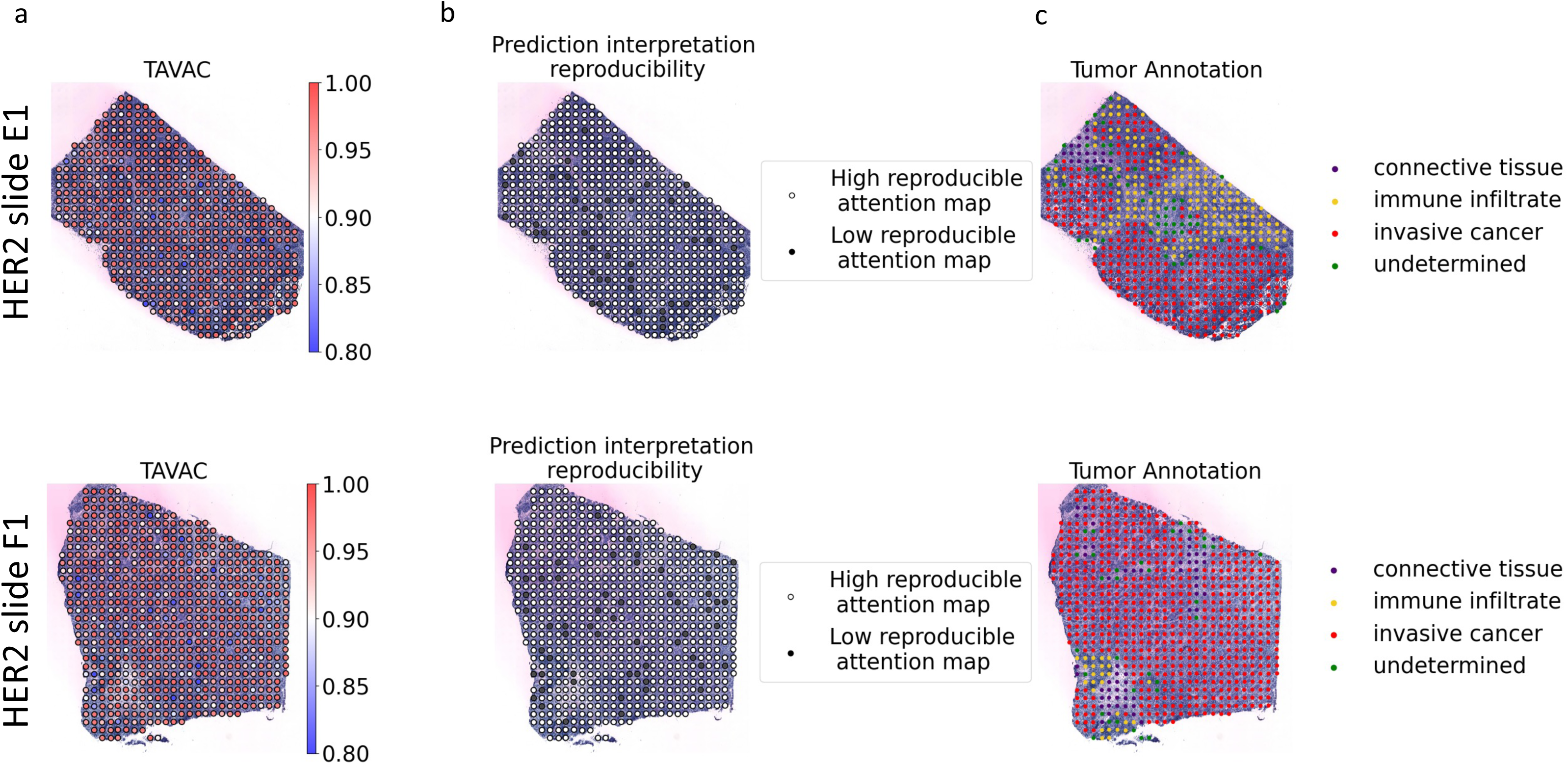
Application of TAVAC scores to filter low-quality fine-grained model interpretation. (**a**) Spots to be excluded (TAVAC < 0.9) are highlighted as (**b**) black dots in the second column due to low TAVAC scores, i.e., their lack of consistency between training and validation attention maps. (**c**) Notably, low TAVAC spots are distributed throughout the tissue, without a distinct correlation to specific pathologist-annotated categories.

## Discussion

TAVAC quantifies overfitting impact for ViT model interpretation. It evaluates the level of generalization in visual features that the models focus on when predicting object types or tumor status at pixel-resolution. TAVAC generalizes to four various types of image benchmark datasets. The algorithm also contributes to the quality assessment of model interpretation, when the overfitting ViT models pays attention to the regions of the images that are away from the object of interest by assigning a lower TAVAC score. Moreover, TAVAC quantifies the level of generalization of histological features that cause ViT models to predict tumor status in a breast cancer H&E image dataset. Moreover, TAVAC is robust on the independent HER2+ breast cancer data. It identifies fine-grained spatial variations of generalization level of model interpretation within the tumor or the normal tissue, respectively, using the pathologist-defined tumor and normal annotations, e.g., connective tissues or immune infiltrate.

Recently, deep learning models focusing on more detailed labeling of smaller cellular clusters (multi-cell) to predict spatial transcriptome data using histological image data (He, et al., 2020; Andersson, et al., 2021) started to uncover complex patterns in the relationship between tissue structure and gene expression, providing new insights into biological processes and diseases. TAVAC measures the level of generalization of deep learning model interpretation, not only on slide-level but also fine-grained level for small groups of cells. The application of TAVAC will allow for robust model interpretation in deep learning models leveraging commonly available H&E-stained images—especially those digitized through modern digital pathology techniques— for high-resolution tumor status or subtype prediction.

Overfitting affects model interpretability (Ramampiandra & Chollet, 2023). TAVAC is the first algorithm focused on the generalization of deep learning model interpretation. Using the TAVAC to filter the low-quality data points where prediction is made based on random noise, increase the confidence in the model interpretation. Future work includes a direct extension of TAVAC for general transformer models (for instance, Natural Language Processing models) where we can extract the attention from the input sequence data. The potential for employing alternative consistency metrics in future explorations remains a viable consideration. Also, optimization and training strategies can be developed to achieve high TAVAC score models. Thus, the TAVAC score can also serve as an independent hyperparameter tuning metric to ensure the generalization of transformer models.

## Methods

### Training Attention and Validation Attention Consistency (TAVAC)

The computation of the Training and Validation Attention Consistency (TAVAC) score is delineated as follows: Commencing with a bifurcation of dataset D into two equal subsets, akin to the methodology of K-fold cross-validation, we designate one subset for training and the other for validation purposes. The initial phase involves training a model on the training subset and appraising its efficacy on the validation subset. Subsequently, we invert the roles of the subsets for a second evaluative iteration. The model’s capacity for accurate prediction and its consistency are gauged by comparing the attention maps—generated via Attention Rollout—for congruent images from the first training phase to those from the second validation phase. These attention maps are classified as either Training Attention or Validation Attention based on their use case. The TAVAC score is then derived by employing Pearson Correlation to assess the concordance between the two types of attention maps, a metric that may be expanded to encompass alternative consistency measures in future research endeavors. The pseudo code of the process is attached in Algorithm.1.

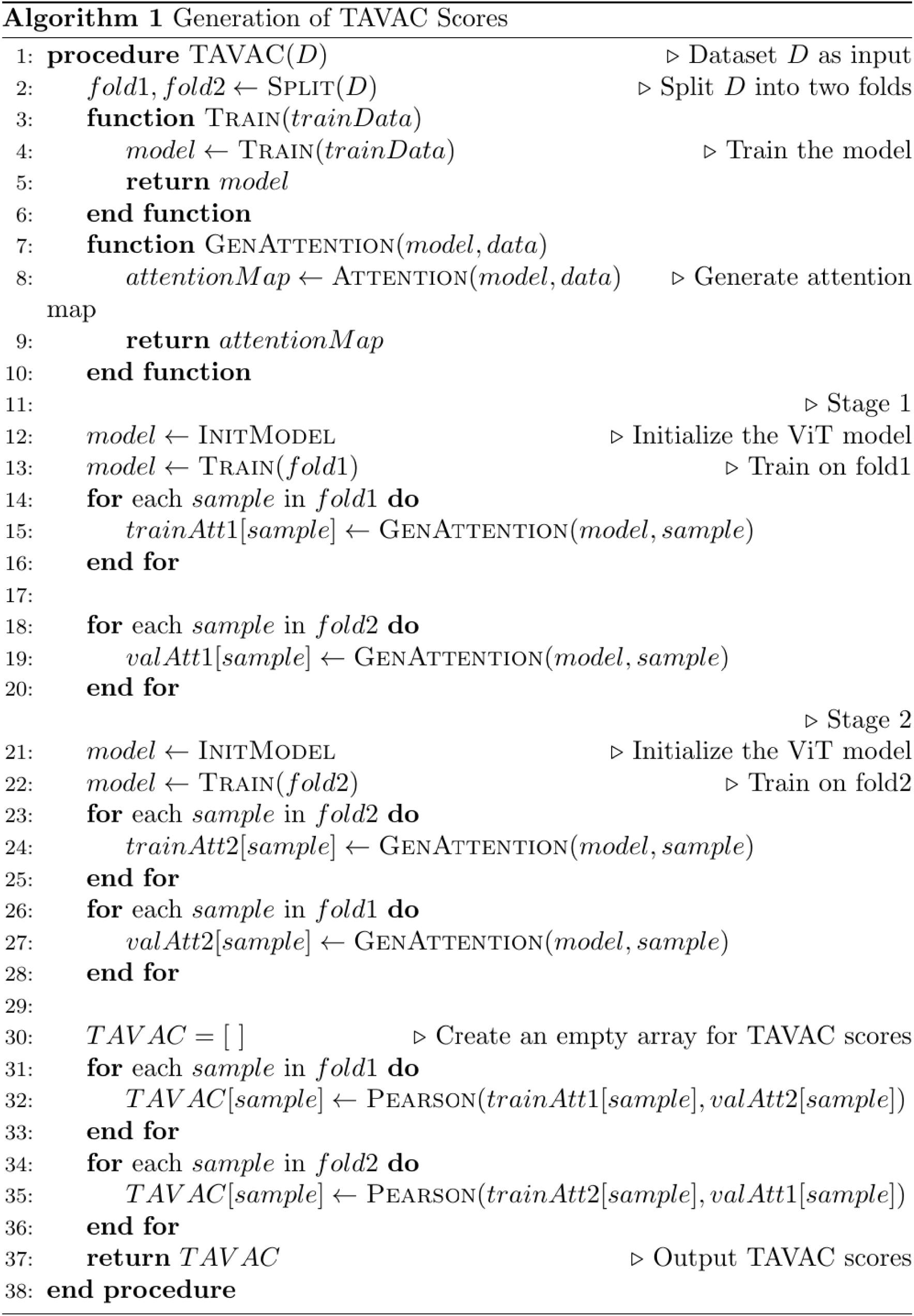

#### Attention Rollout

Attention maps for the ViT models were generated as follows. Attention rollout was used to calculate the attention weight assigned to each pixel of every image. The central architecture of the Vision Transformer model consists of self-attention layers, which allows it to capture long-range spatial relationships within our images (Chefer, et al., 2021). As each pixel or token progresses through the layers of the ViT model, the attention weights of each token are propagated and adjusted accordingly (Abnar & Zuidema, 2020). This enabled us to see how much attention each pixel of the images contributed to the prediction power of the Vit Model.

#### Evaluate TAVAC using scatterplot and histogram

We first use the TAVAC method to evaluate individual images selected from each of our datasets. To this end, both stages of training and validation attention were converted to RGB images, where each pixel of each image was represented by red, green, or blue colors. Thereafter the RGB image of each attention map was converted into a NumPy array, which stored respective attention values. Using the attention values stored in the NumPy arrays, we generated a scatter plot to visualize the train and validation attention correlation for each image. The Pearson correlation coefficient is used to represent the TAVAC score. TAVAC scores close to 1 meant the correlation between train and validation attention values for the particular image was strong, while TAVAC scores close to 0 demonstrated a weak correlation between train and validation attention.

The above allowed us to evaluate the TAVAC for individually selected images. A more holistic approach for overall model interpretation is TAVAC score distribution, represented by a histogram. We calculated the TAVAC score for every image in our respective datasets using a similar procedure to the above. Next, we plotted these values on a histogram to visualize the distribution in attention consistency for the entirety of our datasets. We use mean, median, and variance to compare model performance of good-fit and overfit for all datasets.

#### Pretrained Vision Transformer on ImageNet

For this study, we utilized a particular Vision transformer model pre-trained on Imagenet-21k, an extensive database of 14 million images and 21,843 classes at a resolution of 224×224 (Dosovitskiy, et al., 2020). Images are fed into the transformer encoder as a sequence of equivalently sized linearly embedded patches (16 × 16 resolution) after the addition of classification task (CLS) tokens and positional embeddings. Pretraining the model allows the algorithm to learn to extract meaningful features from images, and this information is then transferred to smaller downstream tasks. In our case, the transfer learning concept enabled the ViT model to employ the knowledge learned on the large ImageNet dataset to our smaller spatial omics breast tissue data (Deng, et al., 2009; Wu, et al., 2020).

## Acknowledgements

Dylan Agyemang’s Summer Student Fellowship was supported by The National Cancer Institute (Award R25CA233420) and The Jackson Laboratory’s Summer Student Program fund. Dr. Sheng Li is supported by the following grants: R35GM133562 (SL), U01HG013175, U01CA271830. Dr. Matthew Mahoney is supported by R01GM141309. Drs. Yue Zhao, Matthew Mahoney, and Sheng Li are supported by 5U54AG079753. We would also like to express our sincere gratitude to senior scientific writer Carmen Robinett, whose invaluable contributions significantly enhanced the quality of this manuscript.

## Code and Data Availability

The code is available in github: https://github.com/LabShengLi/TAVAC.

CIFAR-10, MNIST, FOOD101, CAT AND DOG data are available on hugging face data hub: https://huggingface.co/datasets.

ST-net data is downloaded at: https://data.mendeley.com/datasets/29ntw7sh4r/5

HER2 data is available at: https://github.com/almaan/her2st

## Notes

### Competing Interest Statement

The authors have declared no competing interest.

