## Supplemental Methods and Figures for "Quantifying Interpretation Reproducibility in Vision Transformer Models with TAVAC"

**TAVAC demonstration on individual images within MNIST, FOOD101, and Cats vs. Dogs benchmark datasets**

Similar to the procedure we used to calculate the TAVAC for the horse image in the CIFA10 dataset, we also evaluated the TAVAC for individual images within the other three benchmark datasets: MNIST, FOOD101, Cats vs. Dogs. For each dataset we compare the TAVAC of good-fit and overfitted model for our selected image. This approach is significant in that it allows us to analyze the attention consistency of particular images within each of our datasets instead of the comprehensive distribution of the entire collection of images. For the MNIST dataset, we randomly selected the number 8 image and evaluated the TAVAC by analyzing correlation of train and validation attention. The good-fitted model (Fig. 3) visually shows a near perfect correlation between train and validation attention. The points closely follow the y = x line, culminating in a TAVAC score of 0.99. The overfitted case yields a TAVAC score of 0.98, indicating that the attention consistency between the well-fitted and overfitted models on the 8 images is nearly identical. This suggests that our overfitted case was slightly overfitted. Similarly, we conducted the same process on the FOOD101 data. The well-fitted model exhibited a TAVAC score of 0.97 (Fig. 3). The overfitted model also presents strong attention consistency with a TAVAC score of 0.93 (Fig. 3). Again, the closeness of the two TAVAC scores for the FOOD101 ramen image corresponds to the fact that there was only a marginal difference between the accuracies of the well-fitted and overfitted epoch settings.

Lastly, we evaluated the TAVAC for a dog image in the Cats vs. Dogs dataset for our two model variations. As shown in Fig. 3, along with a TAVAC score of 0.93, there is a strong overall correlation between training and validation attention for the well-fitted model. In contrast, the scatterplot for the overfitted model shows a slightly lower TAVAC score of 0.89. Again, we observe a decrease in the TAVAC score from good-fit to overfitted. Although the discrepancies between good-fit and overfitted cases for the MNIST, FOOD101, and Cats vs. Dogs datasets were marginal, there still exists an inverse relationship between overfitting and TAVAC scores across all three of these benchmark datasets.

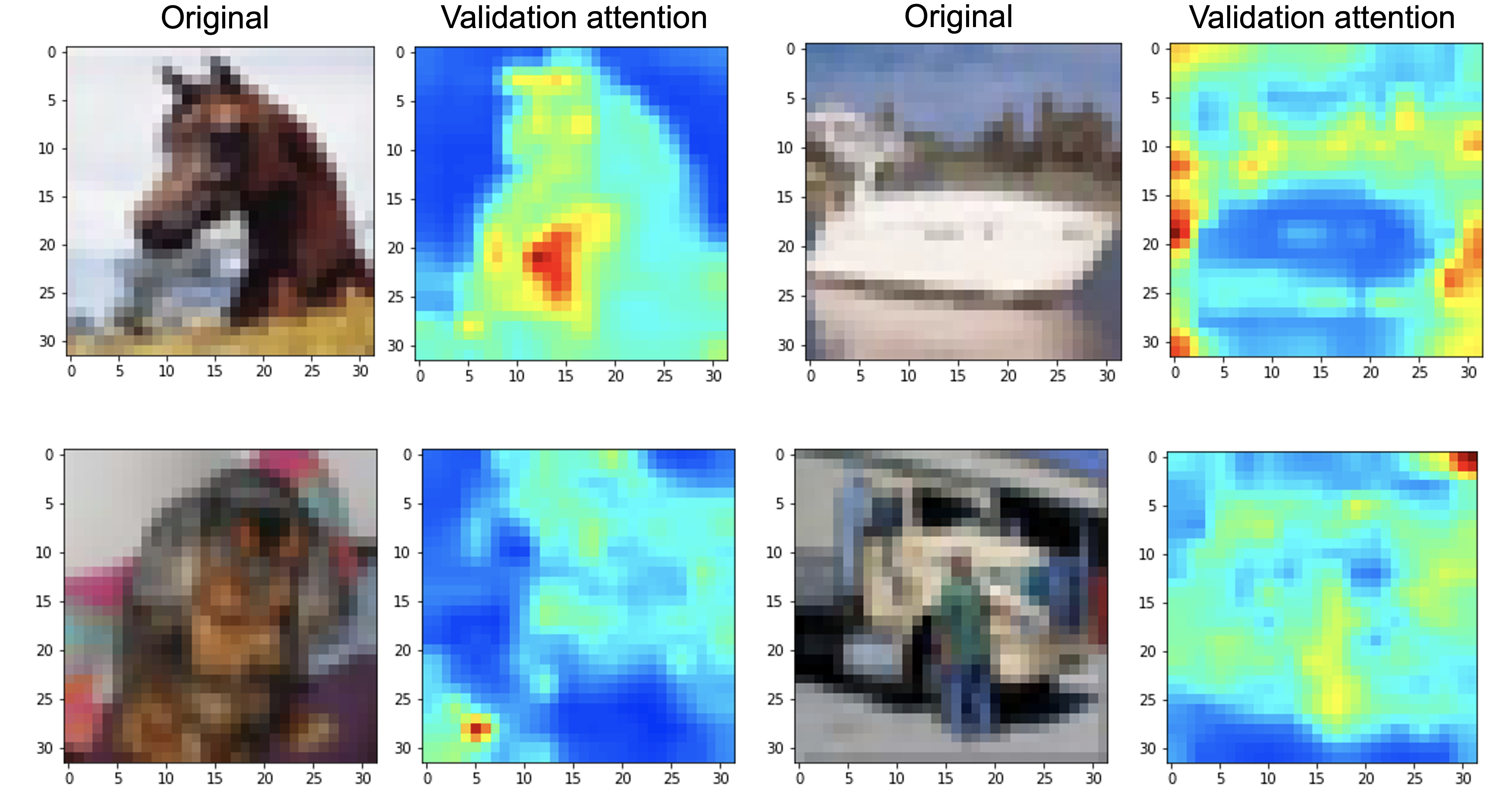

**Figure S1. Examples of manually annotated bad-focus image attention maps for TAVAC evaluation in identifying meaningful high-attention regions.** In ‘bad focus’ group’, the high-attention regions of the validation attention of the *overfitted* model do not overlap with the object of interest.

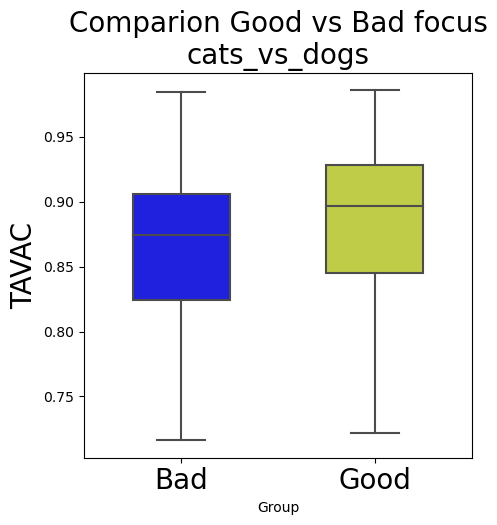

**Figure S2. TAVAC score comparison for images with manually annotated 'Good Focus' versus 'Bad Focus'.** 'Good Focus': manually annotated images where high attention is maintained on the object of interest during validation in well-fitted cases; 'Bad Focus': manually annotated images with poor attention focus during validation in overfitted cases.

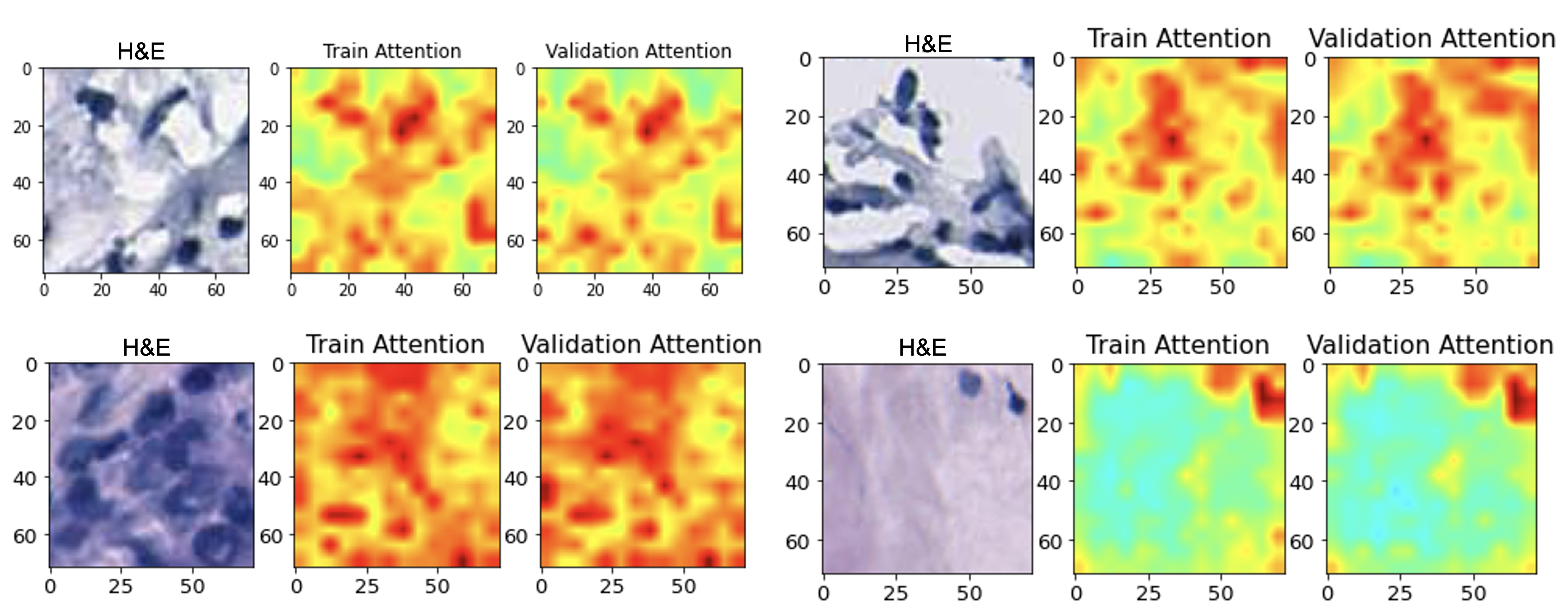
 **Figure S3. High-TAVAC-score attention maps.** Attention maps were derived from ViT models trained with HER2 breast cancer dataset (Andersson, et al., 2021).

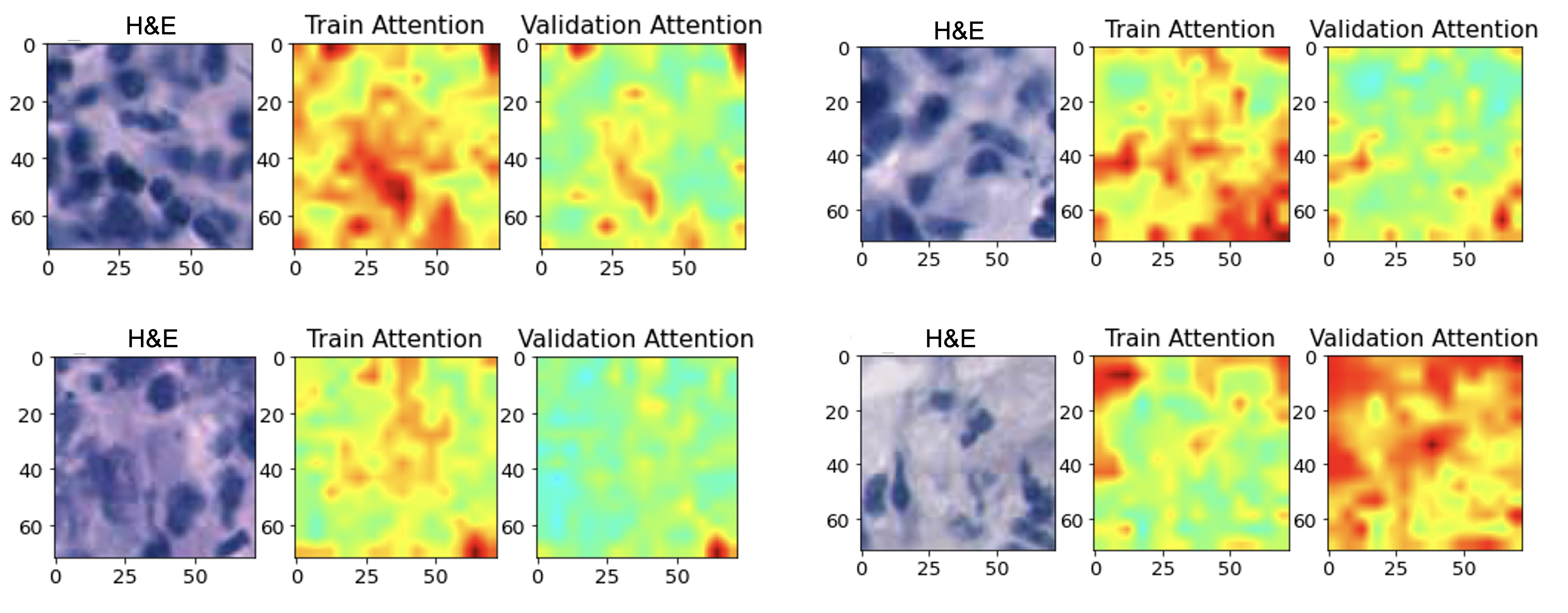

**Figure S4. Low-TAVAC-score attention maps.** Attention maps were derived from ViT models trained with HER2 breast cancer dataset (Andersson, et al., 2021).

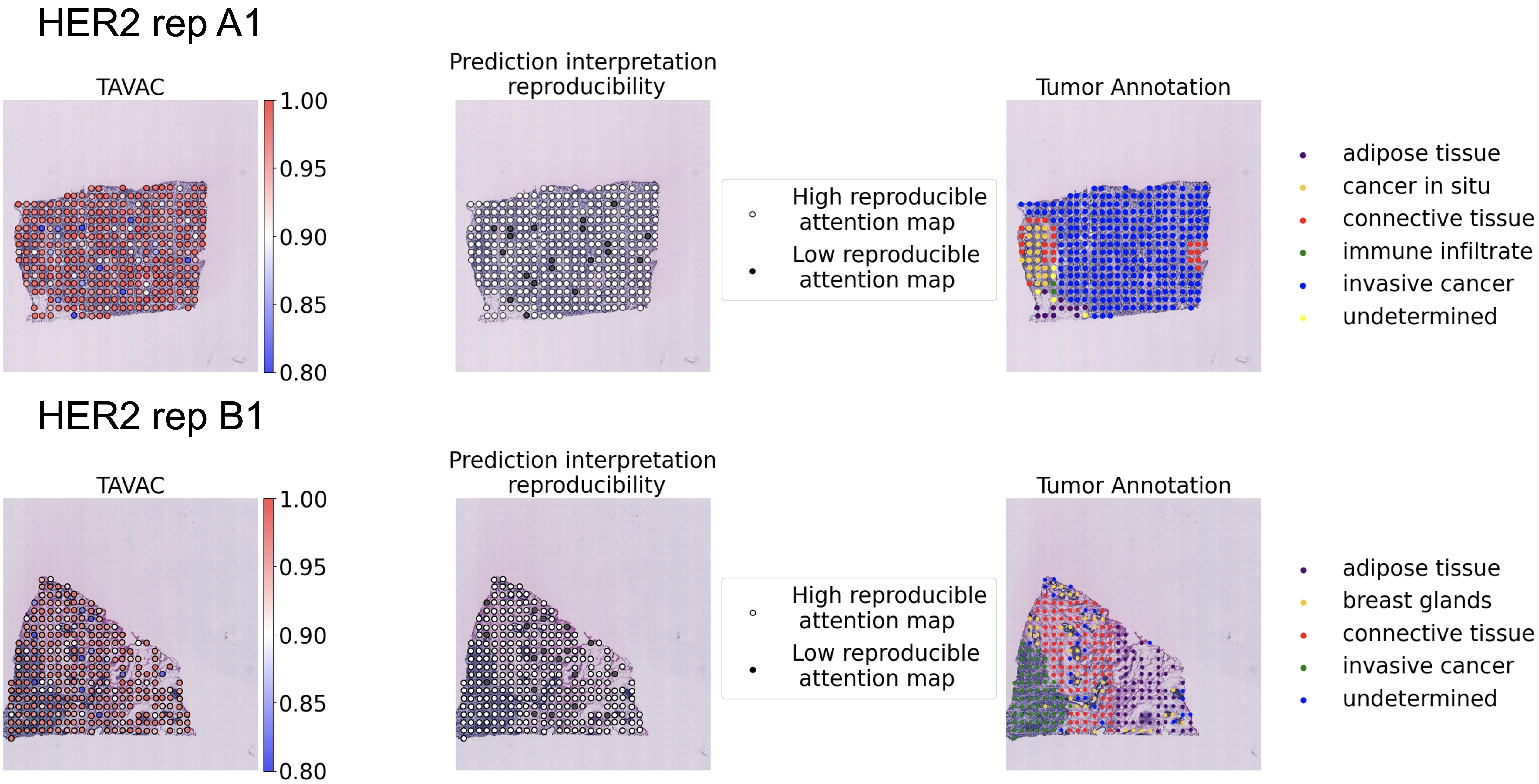

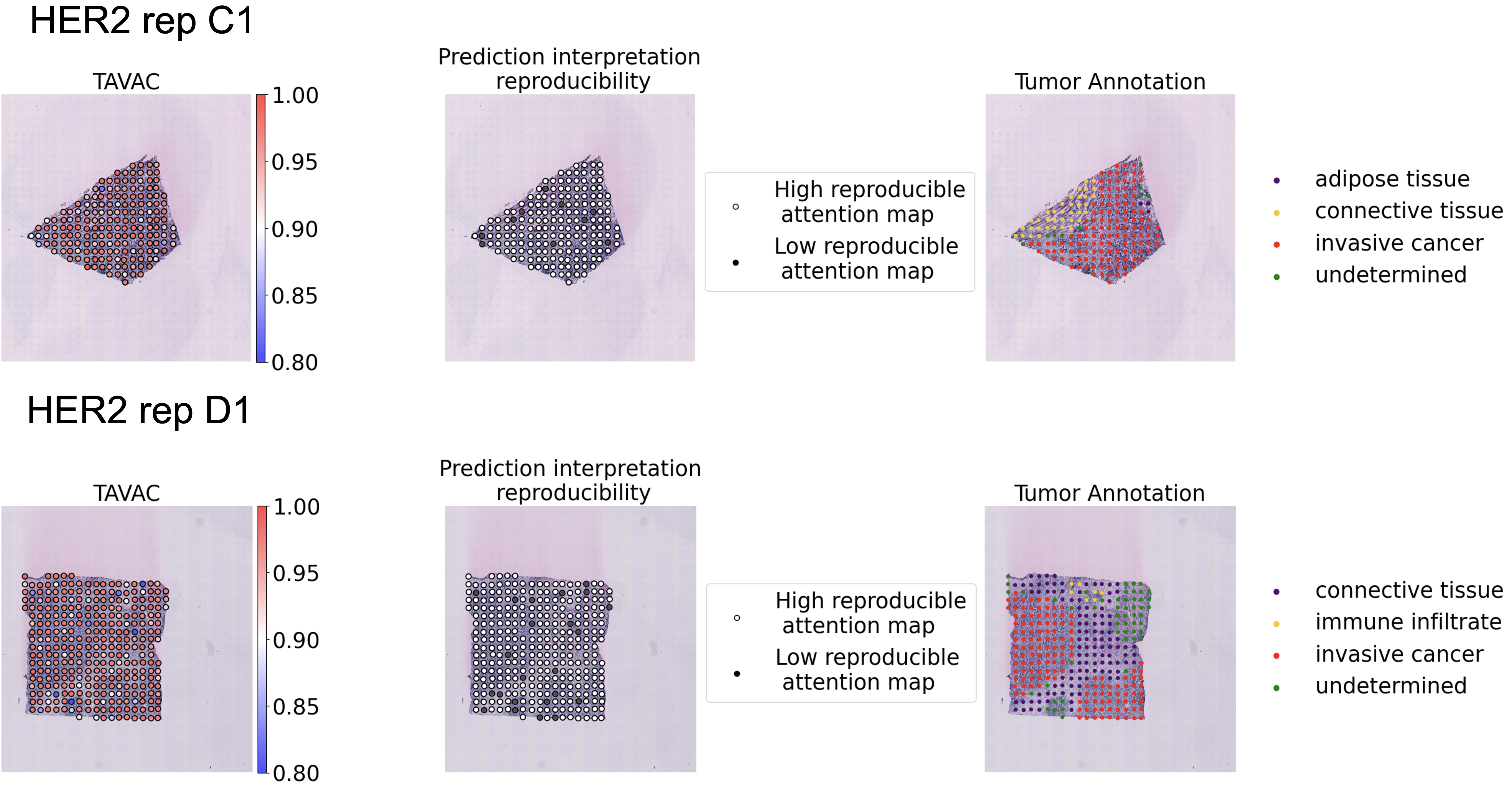
**Figure S5. Application of TAVAC scores to filter low-quality fine-grained model interpretation.** More attention maps were derived from ViT models trained with HER2+ breast cancer dataset (Andersson, et al., 2021). Spots to be excluded (TAVAC < 0.9) are highlighted as black dots in the second column due to low TAVAC scores, i.e., their lack of consistency between training and validation attention maps. Notably, low TAVAC spots are distributed throughout the tissue, without a distinct correlation to specific pathologist-annotated categories.
